# Exosomes from Nef expressing monocytic cells restrict HIV-1 replication in infected cells through the assembly of stress granules

**DOI:** 10.1101/148791

**Authors:** Mohammad Yunus Ansari, Hasan Imam, Nishant Kumar, Zulfazal Ahmed, Shahid Jameel

## Abstract

Exosomes are membranous vesicles secreted from almost all types of cells, carry proteins and nucleic acids and function as vehicles for intercellular communication. Cells infected with HIV-1 or expressing the viral Nef protein secrete more exosomes than uninfected cells or those not expressing this protein. We used stably transfected, Nef-expressing U937 human monocytic cells and exosomes purified from these cells to study their effects on HIV-1 infected and uninfected CD4+ T-cells. The Nef exosomes inhibited virus production from HIV-1 infected CD4+ T-cells, but caused activation induced cell death in uninfected bystander cells. Mutations in its conserved Arginine residues and in the secretion-modification-region failed to secrete Nef into exosomes. Cell lines expressing these mutant Nef proteins did not deliver it to the target CD4+ T-cells, and exosomes prepared from these mutant Nef-expressing cells also did not inhibit virus production. Nef exosomes inhibited virus production by inducing the assembly of stress granules in HIV-1 infected cells, which sequestered increased amounts of gag mRNA. This is a novel mechanism wherein we show the effects of exosomes on the assembly of stress granules and viral translational repression.

## Introduction

The human immunodeficiency virus (HIV) continues to be a major public health threat, with an estimated 35 million people living with the infection [1]. The development of antiretroviral drugs is a major advance, which has improved the duration as well as quality of life following HIV infection. Although therapeutic use of these drugs can reduce the viral load in infected individuals to below detection levels, they cannot clear the infection. Viral latency poses the biggest challenge to HIV eradication [2]. During the latent phase, HIV remains integrated in the genome of infected cells, but does not replicate actively and can therefore stay hidden from the host immune system. Multiple signals, which are not fully understood, or discontinuation of therapy may result in active transcription and viral replication [3-5]. The mechanisms of latency include transcriptional regulation of viral replication and microRNA (miRNA)-mediated posttranscriptional regulation of viral mRNA translation and/or degradation [5-7].

Exosomes are 30-100 nm vesicles produced by the inward invagination of endosome membranes in multivesicular bodies (MVBs), which fuse with the plasma membrane to release exosomes into the extracellular space [8, 9]. These are secreted from almost all cell types and are present in many biological fluids including blood, saliva, breast milk, urine and semen [10-13], and have been proposed as biomarkers for several types of cancers [10, 14] kidney dysfunction [15] and liver disease [16]. Exosomes contain many biologically active molecules including proteins, messenger RNAs (mRNAs), miRNAs and lipids, and are suggested to be important for cell-to-cell communication [17-19]. For example, alpha interferon (IFN-α) treated nonparenchymal liver cells, which are resistant to hepatitis B virus (HBV) secrete exosomes that can transfer virus resistance to HBV permissive hepatocytes [20]. The tumor suppressor protein PTEN is exported through exosomes and can perform its function in target cells [21]. Exosomes produced from HIV-1 infected cells contain the transactivation response (TAR) element and exposure of naïve cells to these exosomes increases their susceptibility to HIV-1 infection [22]. The HIV-1 genome is also secreted into exosomes from infected cells, and the region responsible for this transfer was located at the 5’ end of the p17^Gag^ open reading frame. The exosomal secretion of miR29 in HIV-1 infection increases upon opiate abuse; this reduces the expression of platelet-derived growth factor-beta in recipient cells and plays an important role in HIV-associated neurological disorders[23].

The HIV-1 genome encodes the prototypic retroviral proteins (Gag, Pol and Env), two regulatory proteins (Tat and Rev) and four accessory proteins (Nef, Vif, Vpr and Vpu). Of these, the Nef protein is expressed early in the viral life cycle and is secreted into exosomes, which are found in the blood circulation of infected individuals [24]. Motifs in the Nef protein that are required for its packaging into exosomes have been mapped to a stretch of four arginine residues (amino acids 17-22) and the sequence VGFPV (amino acids 66-70) named as the secretion modification region (SMR) [25]. Earlier work from our laboratory has shown the Nef protein expressed to interact with the miRNA silencing machinery in human monocytic U937 cells and to be packaged into exosomes [26]. It also alters the mRNA and miRNA profile of exosomes [27]. While exosomes containing Nef were shown to cause the activation-induced death of bystander CD4+ T cells [28], their effects on HIV-1 infected cells have not been explored. We show here that Nef-containing exosomes inhibit viral replication in infected CD4+ T cell lines through a post-transcriptional mechanism that involves stress granule formation and translational suppression of viral mRNAs. This further emphasizes the dual effects of Nef in optimizing the cellular environment for viral replication and persistence.

## Materials and Methods

### Cells

The J1.1 and U1 cells were obtained from NIH AIDS Reagent Bank and maintained in RPMI containing 10% fetal bovine serum (FBS). The U937/Nef-EYFP and U937/EYFP stable cell lines were reported previously [26] and were maintained in RPMI containing 10% FBS and 350 ng/ml Puromycin. The HEK293T and TZM-bl cells were maintained in DMEM containing 10% FBS. The U937/Nef(4R/4A)-EYFP and U937/Nef(VGFPV)-EYFP stable cell lines were generated in the laboratory. For this, mutations within the *nef* gene were first generated in the pMSCV-Nef-EYFP plasmid background [24] at Mutagenex Inc, NJ, USA. Retroviruses expressing Nef(4R/4A)-EYFP and Nef(VGFPV)-EYFP were generated by cotransfection of HEK293T cells with 2 μg of the transfer plasmid, 1 μg of pGag-Pol and 0.5 μg of pVSVg in a T25 flask using the calcium phosphate method. The culture supernatants were collected after 36 hr and used as the source of recombinant retroviruses. Human monocytic U937 cells were washed with RPMI, starved for 90 min without serum and then transduced with 500 μl of culture supernatants per 1×10^6^ cells. After a 4 hr adsorption step, the cells were washed and kept in complete medium for 48 hr prior to the addition of 350 ng/ml Puromycin. The cells were split every 48 hr and those surviving after 5 passages were used for the analysis. The clones were sorted for the EYFP positive population using a Becton Dickinson Aria Cell Sorter in the Central Facility of the National Institute of Immunology, New Delhi, India. The sorted clones were cultured for 4-5 passages and checked for purity and EYFP expression using a Cyan-ADP flow cytometer (Beckman Coulter). Data was analyzed using Summit 4.3 software. Characterization of Nef function in the U937/Nef(4R/4A)-EYFP and U937/Nef(VGFPV)-EYFP stable cell lines was done by flow cytometry for the surface expression of CD4, MHC I, CD80 and CD86, with CD54 as a negative control (data not shown).

### Antibody and plasmid constructs

Murine anti-p24 monoclonal antibody (hybridoma supernatant) was obtained from NIH AIDS Reagent Bank. APC or PE labeled antibodies for CD80 (13-0809), CD81 (17-0819), CD54 (13-0549) and MHC-1 (17-9876) were obtained from eBiosciences. Anti-GFP (6556) antibody was purchased from Abcam. Antibodies for phosphor-eIF2α (3597) and eIF2α (9722) were from Cell Signaling, and those for G3BP1 (HPA004052) and APOBEC3G (HPA001812) were from Sigma. The G3BP1-GFP expression plasmid was a kind gift from Dr. Jomon Joseph [29]. The p24 ELISA kit was obtained from the NIH AIDS Reagent Bank.

### Transfection

The HEK293T and TZM-bl cells were transfected using the JetPrime transfection reagent according to the manufacturer’s protocol. The cells were seeded in 12-well plates and transfected with 200 ng of the G3BP1-GFP plasmid. For transfection of J1.1 cells a Nucleofection kit (Lonza) was used according to the manufacturer’s protocol.

### Culture and analysis of J1.1 cells

For the coculture of J1.1 cells with U937/EYFP, U937/Nef-EYFP or Nef mutant cell lines, 0.5×10^6^ cells/ml each of J1.1 cells and the U937 stable cell line were mixed in 5 ml RPMI containing 10% FBS. Supernatants were harvested from the mixed culture at different time points and analyzed for p24 levels by Western blot or ELISA, and the cells were analyzed by flow cytometry. For preparing conditioned media, U937/EYFP and various U937/Nef-EYFP stable cell lines were seeded at a density of 0.5×10^6^ cells/ml in RPMI containing 10% FBS for 72hr. Cultures were centrifuged at 1,000 xg for 10 min to remove cells and the cell-free media were diluted 1:1 with fresh RPMI containing 10% FBS. These were used as conditioned media to grow J1.1 cells.

### Purification and characterization of exosomes

Exosomes were purified from culture supernatants by multiple rounds of centrifugation as described earlier [26]. In brief, the U937 stable cell lines were cultured in RPMI containing 10% FBS from which contaminating exosomes and microvesicles had been removed by centrifugation as described earlier [24]. The cells were cultured at a density of 0.5×10^6^ cells/ml for 72 hours. These were removed by centrifugation at 1,000 xg for 10 min, and the supernatants (S1) were then centrifuged at 15,000 xg to remove cell debris and other aggregates. The clarified supernatants were then centrifuged at 100,000 xg to pellet exosomes. The exosomal pellet was washed with ice cold PBS and resuspended in 100 μl PBS for further analysis by flow cytometry and Western blotting. To purify exosomes for live cell experiments, the ultracentrifugation step was substituted by ultrafiltration. Briefly, 15 ml of the culture supernatants from EYFP or Nef-EYFP cell lines were concentrated through 100,000 MWCO amicon filters (Millipore) to a final volume of 200 μl. Flow cytometric analysis of exosomes was carried out as described elsewhere [30]. Briefly, 10 μg of purified exosomes were coated on 4 μm aldehyde/sulfate latex beads (Invitrogen) in PBS overnight at 4°C followed by blocking with 100 mM glycine. The coated beads were washed with PBS/0.5% BSA and resuspended in 0.5 ml of the same. From this, 10 μl of the coated beads were used for flow cytometric analysis of exosomal protein markers.

### Fluorescence imaging

The formation of stress granules was analyzed by confocal microscopy. Cells (J1.1, HeLa or HEK293T) were transfected with the G3BP1-GFP expression plasmid. After 24 hr, the transfected cells were treated with purified exosomes or conditioned media from control or Nef-expressing U937 cells. Following 48 hr of exposure, the cells were washed with RPMI, fixed in 4% paraformaldehyde and mounted using antifade containing DAPI (Invitrogen, Carlsbad, CA, USA). Images were acquired using a Nikon A1/R confocal microscope at 60X magnification.

## Results

### Exosomal Nef reduces HIV-1 replication in infected CD4+ T cell line

To evaluate the effects of exosomes produced from Nef-expressing monocytic cells on HIV-1 infected CD4+ T cells, we have used U937 cells stably expressing the Nef-EYFP fusion protein (or EYFP as a control) [26], and J1.1 cells, which are Jurkat cells latently infected with HIV-1. We first evaluated virus production from J1.1 cells as a function of time when these cells were co-cultured with U937/Nef-EYFP or U937/EYFP cells and. Equal volumes of the culture supernatants were collected at each time and analyzed for HIV-1 in the culture medium by Western blotting for p24. The result showed lower levels of HIV-1 from J1.1 cells that were cocultured with U937/Nef-EYFP compared to U937/EYFP cells (Fig.1A). To confirm this was not due to activation of J1.1 cells by the control U937/EYFP cells and increased virus production, we also cultured J1.1 cells alone. Equal volumes of the supernatants were collected at each time and analyzed by Western blotting for p24. We detected similar levels of virus in the culture supernatants of J1.1 cells alone or J1.1 cells co-cultured with U937/EYFP and decreased virus production in case of U937/Nef-EYFP co-culture (Fig. S1). The levels of virus secreted in the culture media were quantitatively estimated using a p24 ELISA. There was around 50% reduction in virus production from J1.1 cells co-cultured with U937/Nef-EYFP compared to U937/EYFP cells (Fig. 1B). To learn if this effect was due to physical contacts between the U937 and J1.1 cells or due to secretory factor(s), we used cell-free conditioned media from U937/Nef-EYFP or U937/EYFP cells to culture J1.1 cells. There was reduced intracellular viral replication and secretion in J1.1 cells treated with conditioned media from U937/Nef-EYFP compared to U937/EYFP cells (Fig. 1C and Fig. S1), suggesting that secretory factor(s) might mediate this effect. Conditioned media from U937/Nef-EYFP cells had no significant effects on the viability (Fig. S2A) or proliferation (Fig. S2B) of J1.1 cells. Similar results were also obtained when U1 cells, which are U937 cells that are latently infected with HIV-1, were either co-cultured or treated with conditioned media from U937/Nef-EYFP cells (Fig. S3).

**Figure 1:**
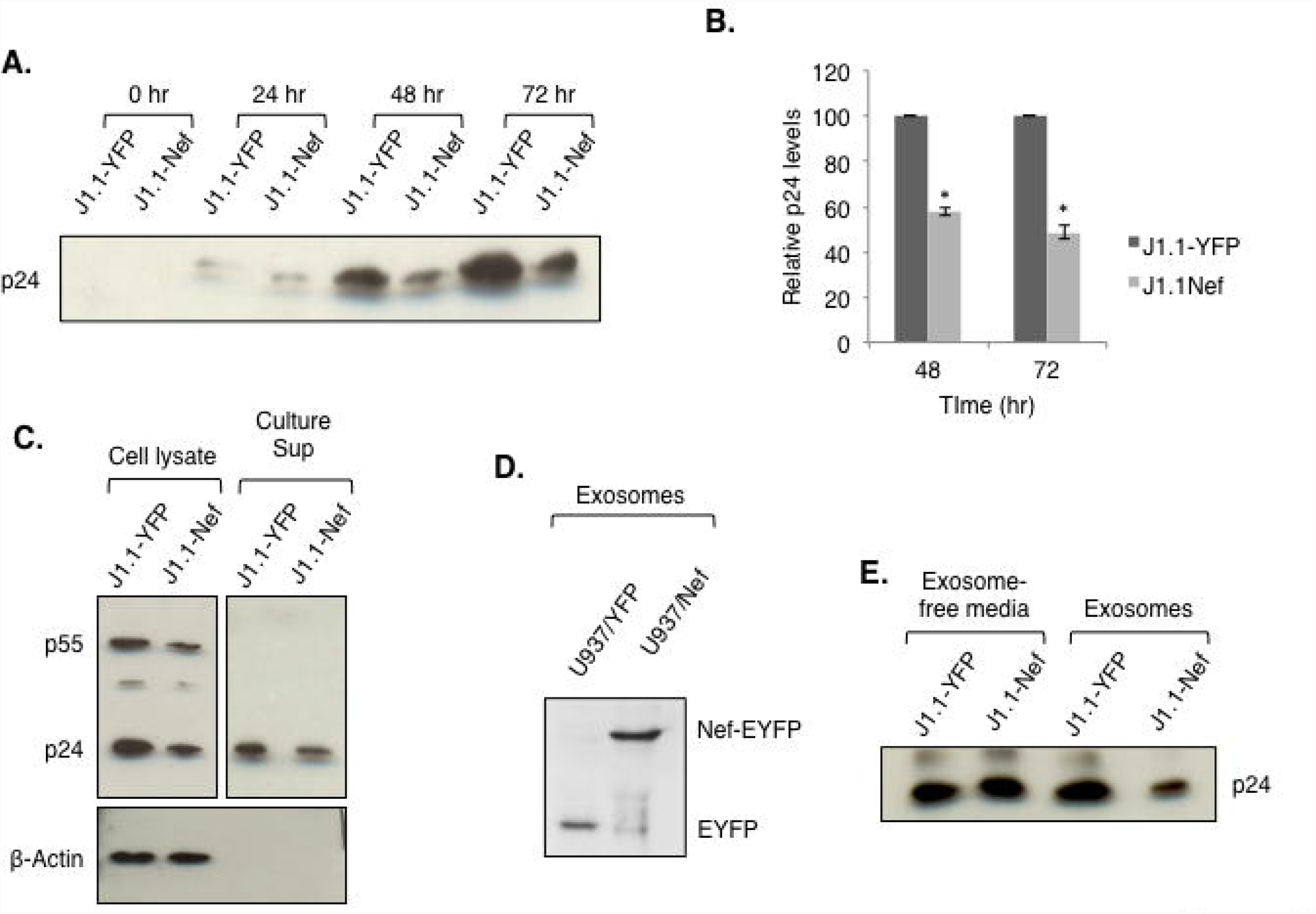
Exosomes from Nef-expressing cells inhibit virus production from infected cells. (A) J1.1 cells were co-cultured with U937/EYFP cells (lanes J1.1-YFP) or U937/Nef-EYFP cells (lanes J1.1-Nef). The culture supernatants were collected at different times and analyzed by Western blotting for p24. (B) The coculture supernatants from J1.1 and either U937/Nef-EYFP cells (J1.1-Nef) or U937/EYFP cells (lanes J1.1-YFP) were quantified with ELISA for p24 levels. Results are shown as mean of three independent experiments, each with triplicate measurements (*p<0.05). (C) J1.1 cells were cultured in conditioned media from U937/EYFP cells (lanes J1.1-YFP) or U937/Nef-EYFP cells (lanes J1.1-Nef) for 72 hr. Cells and supernatants were harvested and analyzed for intracellular p24/p55 and p24, respectively. Beta-actin served as a loading control for cell lysates. (D) Exosomes were purified from U937/EYFP or U937/Nef-EYFP cells and Western blotting was carried out with anti-GFP antibodies; the EYFP and Nef-EYFP are indicated. (E) J1.1 cells were treated with exosome-free media or exosomes from U937/EYFP cells (lanes J1.1-YFP) or U937/Nef-EYFP cells (lanes J1.1-Nef) for 72 hr, and p24 in the culture supernatants was analysed by Western blotting.

The effects of conditioned media suggested that secretory factor(s) from Nef-expressing cells mediate the down modulation of HIV-1 production from J1.1 cells. Since Nef is also secreted in exosomes [27], it is plausible that these effects are mediated through exosomes. We therefore isolated exosomes from the culture supernatants of U937/Nef-EYFP and U937/EYFP cells (Fig. S4A), and characterized the purified exosomes for surface markers with flow cytometry (Fig. S4B). Secretion of the Nef protein in exosomes was confirmed by Western blotting (Fig. 1D). Exosomes produced from Nef-expressing CD4^+^ T cells were shown earlier to cause apoptosis in bystander CD4^+^ T cells [28]. To functionally characterize our exosome preparation, we isolated peripheral blood lymphocytes (PBLs) from donor blood, treated these with exosomes from U937/Nef-EYFP or U937/EYFP cells and checked for apoptosis by flow cytometry for Annexin V staining (Fig. S4C) and immunoblotting for PARP cleavage (Fig. S4D). The results showed that exosomes secreted from U937/Nef-EYFP cells induced apoptosis in bystander CD4^+^ T cells. We also tested if these exosomes induced apoptosis in HIV-1 infected J1.1 cells by analyzing the cleavage of procaspase 3 to caspase 3. Surprisingly, no induction of apoptosis was observed in HIV-1 infected cells treated with conditioned media from U937/Nef-EYFP cells (Fig. S5A). The interaction of Nef with CXCR4 is also reported to be important for inducing apoptosis in bystander CD4^+^ T cells [31]. We therefore quantified CXCR4 levels in J1.1 cells and found no difference compared to the parent Jurkat cell line (Fig. S5B).

To determine if Nef exosomes or some other secretory component(s) are responsible for reduced viral replication in J1.1 cells, we treated J1.1 cells with exosomes or the exosome-free (EF) fraction from U937/Nef-EYFP and U937/EYFP culture supernatants. The result showed decreased virus production from J1.1 cells only when these were treated with Nef exosomes but not with the exosome-free fraction (Fig. 1E). These results confirm that exosomes produced from Nef-expressing cells inhibit virus production in infected CD4+ T cells.

### The exosomal delivery of Nef to HIV-1 infected CD4+ T cells is important for inhibiting viral replication

Since Nef is secreted in exosomes, we asked whether the effects on viral replication are linked to the presence of Nef in exosomes. To determine the role of Nef, we generated U937 based cell lines that stably expressed two Nef mutants that are compromised for secretion into exosomes [25]. These include the Nef-4R/4A mutant with arginine to alanine changes at positions 17, 19, 21 and 22, and the Nef-VGFPV mutant in which residues in the SMR region (amino acids 66-70) were changed to alanine. Flow cytometry (data not shown) and Western blotting were used to confirm the expression and exosomal exclusion of these mutant Nef-EYFP fusion proteins (Fig. 2A).

**Figure 2:**
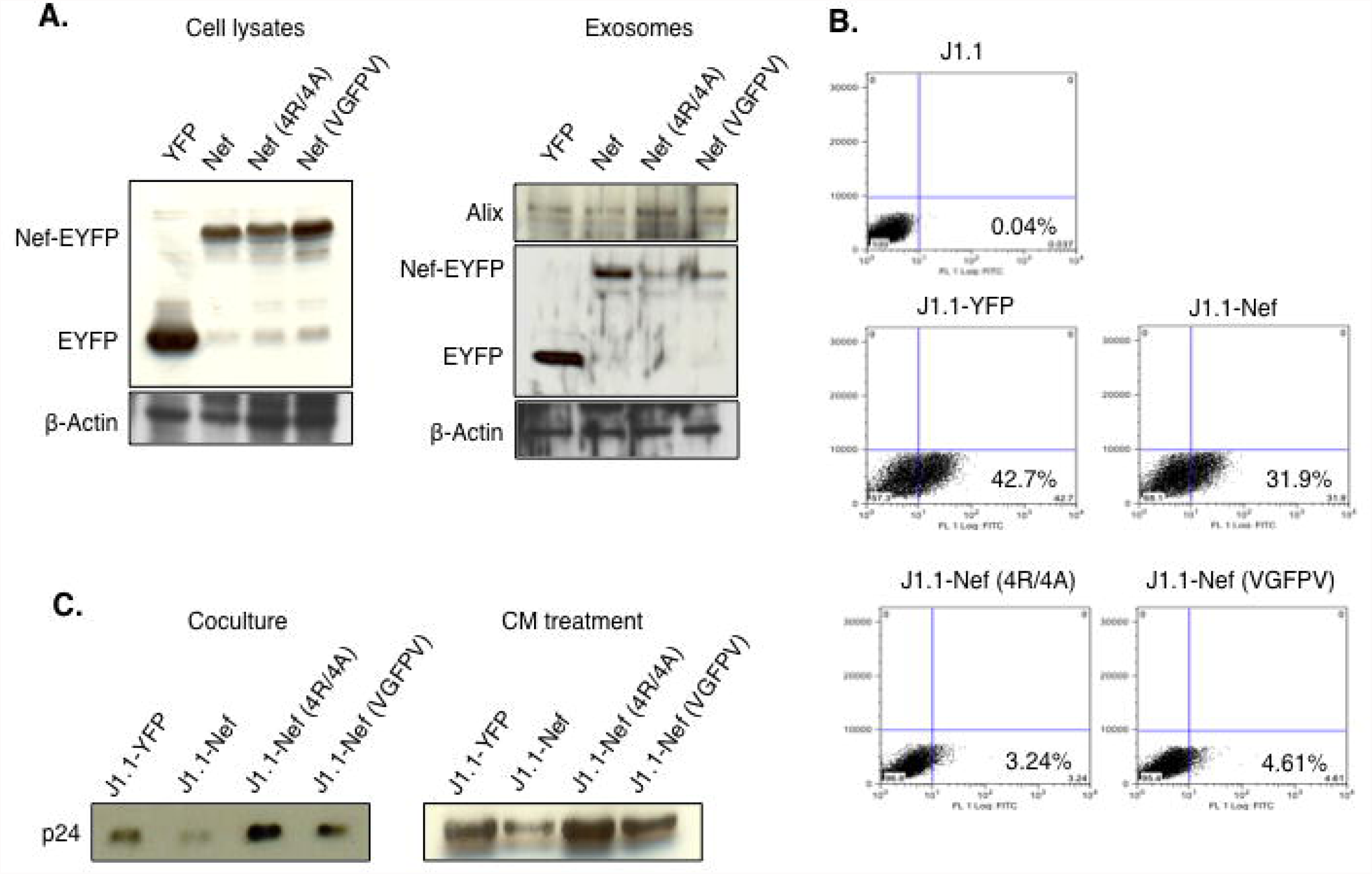
Exosomal secretion of the Nef protein and its delivery into target cells. (a) U937 cell lines that stably expressed the two Nef mutants (4R/4A and VGFPV) were generated. The expression of wild type and mutant Nef proteins in the cell lysates and exosomes purified from these cell lines were evaluated by Western blotting with anti-GFP antibodies; the EYFP and Nef-EYFP are indicated. Beta-actin served as a loading control. Exosomes were also marked by Western blotting for Alix. (B) Delivery of the Nef protein through exosomes was analyzed by coculturing J1.1 cells with U937 cells stably expressing EYFP, Nef and the two Nef mutants followed by flow cytometry for EYFP in J1.1 cells. The J1.1 and U937 cell populations were separated based on forward scatter as shown in Supplementary Figure 6. The numbers indicate percent J1.1 cells positive for EYFP. (C) J1.1 cells were either cocultured with or treated with conditioned media (CM) from U937/EYFP, U937/Nef-EYFP, U937-Nef(4R/4A)-EYFP or U937-Nef(VGFPV)-EYFP cells. The culture supernatants were collected after 72 hr and analyzed for p24 by Western blotting.

Flow cytometry was then used to analyze the delivery of wild type and mutant Nef proteins into J1.1 cells via exosomes. The J1.1 cells were cultured with U937 cells stably expressing Nef-EYFP, Nef4R/4A-EYFP, Nef-VGFPV-EYFP or just EYFP. After fixing, the mixed cell population was gated for the U937 and J1.1 cell populations (Fig. S6), and the latter quantified for the EYFP signal. The results showed about 30-40% of J1.1 cells co-cultured with U937/EYFP or U937/Nef-EYFP cells to be positive for the EYFP signal, but this reduced about 8- to 10-fold for J1.1 cells co-cultured with either U937/Nef4R/4A-EYFP or U937/Nef-VGFPV-EYFP cells (Fig. 2B). These results directly indicate that Nef is efficiently transferred from U937 cells to J1.1 cells through exosomes. We then looked at the effects of mutant Nef proteins on down modulation of viral replication in J1.1 cells. Both Nef mutants that were compromised for secretion into exosomes and transfer into J1.1 cells also did not show reduced HIV-1 production from J1.1 cells, either on co-culture with U937 stable cell lines or when conditioned media from U937 stable cell lines were used to treat J1.1 cells (Fig. 2C).

### Nef exosomes induce markers of translational repression and assembly of stress granules in target cells

To understand the mechanism(s) underlying the effects of Nef exosomes on HIV-1 replication, we first tested if these affect HIV-1 LTR activity. For this, J1.1 cells were transfected with a HIV-1 LTR-luciferase reporter construct, and these were either co-cultured or treated with conditioned media from U937/Nef-EYFP or U937/EYFP cells. After 48 hr, the cells were analyzed for HIV-1 LTR driven luciferase activity. No significant differences in luciferase activities were observed in either condition between Nef and control exosomes (Fig. S7A and S7B). We also quantified gag mRNA levels in J1.1 cells upon treatment with conditioned media or exosomes from U937/Nef-EYFP or U937/EYFP cells and found no significant differences (Fig. S7C and S7D). These observations suggested post-transcriptional regulation of HIV-1 replication by Nef-containing exosomes. To check the role of miRNAs in posttranscriptional regulation of gag mRNA we examined miRNA-mediated silencing of HIV-1 mRNA. For this, J1.1 cells were transfected with plasmid pMIR-Report-Nef3’UTR [32], which carries the HIV-1 nef-3’UTR downstream of the luciferase reporter gene; the plasmid pMIR-Report that lacked the nef-3’UTR was used as a control. The transfected cells were treated with exosomes purified from U937 cells that stably expressed either wild type or mutant Nef-EYFP fusion proteins or EYFP. There were no significant differences in luciferase activities in J1.1 cells treated with any of the exosomes (Fig. S7E).

We then explored the possibility of translational suppression of viral mRNAs. For this, we first checked phosphorylation of the eukaryotic translation initiation factor alpha (eIF2α), which is an indicator of translation arrest [33]. There was increased phosphorylation of eIF2α in J1.1 cells following treatment with exosomes containing Nef compared to control exosomes (Fig. 3A). It is reported that phosphorylation of eIF2α may lead to the assembly of stress granules, which are sites for the translational repression of selected mRNAs [34]. Under such conditions the expression of G3BP1, a marker of stress granules, increases and helps in their assembly. The J1.1 cells treated with Nef exosomes also showed increased levels of G3BP1 compared to cells treated with control exosomes (Fig 3A). Stress granules sequester selective populations of mRNAs leaving out those mRNAs whose products are needed to overcome stress. We did not observe any reduction in the levels of Hsp70, a molecular chaperone that is needed to resolve cellular stress (Fig 3A).

**Figure 3:**
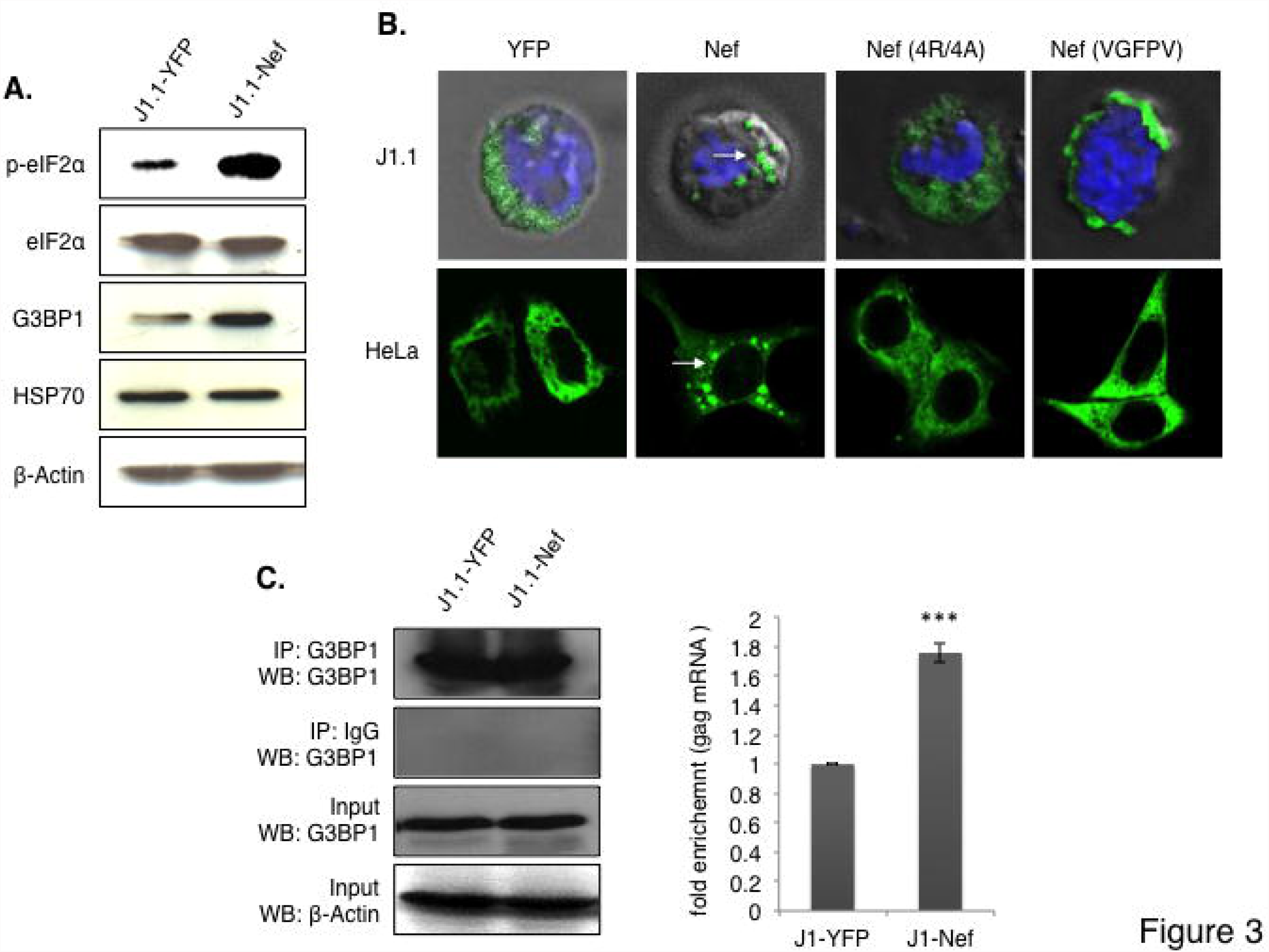
Nef exosomes induce stress granules assembly and sequestration of HIV-1 gag mRNA in J1.1 cells. (A) J1.1 cells were treated with conditioned media from U937/EYFP cells (J1.1-YFP) or U937/Nef-EYFP cells (J1.1-Nef) for 72 hr. The cell lysates were then analyzed for phosphor-eIF2α, total eIF2α, G3BP1 (stress granule marker) and HSP70 levels by Western blotting. Beta-actin served as a loading control. (B) J1.1 cells were transfected with the G3BP1-GFP plasmid and seeded in a 12-well plate, followed by treatment with exosomes purified from U937/EYFP, U937/Nef-EYFP, U937-Nef(4R/4A)-EYFP or U937-Nef(VGFPV)-EYFP cells. Assembly of stress granules (arrows) was checked by confocal microscopy. (C) Stress granules were immunoprecipitated with anti-G3BP1 antibody and association of gag mRNA with stress granules was analyzed by qRT-PCR. Left panel shows Western blot of immunoprecipitation to confirm pull down of G3BP1. Input G3BP1 and actin were used as control. Right panel shows the fold enrichment of gag mRNA in stress granules. RNA was isolated from anti-G3BP1 immunoprecipitate by Trizol-Chloroform and analyzed for gag mRNA by qRT-PCR (***p<0.005).

Furthermore, we tested the assembly of stress granules by first transfecting a G3BP1-GFP reporter construct in J1.1 cells followed by treatment with exosomes from Nef-EYFP, mutant Nef-EYFP and EYFP cell lines. Stress granule assembly was observed in J1.1 cells treated with exosomes from U937/Nef-EYFP cells, but not from control cells or those that express mutant Nef proteins (Fig. 3B and Fig. S8). Similar results were obtained in HeLa cells transfected with the G3BP1-GFP reporter followed by treatment with different exosome populations (Fig 3B). To ascertain if gag mRNA is increasingly associated with stress granules resulting in attenuated translation, we immunoprecipitated stress granule and quantified the levels of gag mRNA in the precipitates. Stress granules were immunoprecipitated with anti-G3BP1 antibodies, total RNA associated with stress granules was isolated and gag RNA was quantified by qRT-PCR. Higher levels of gag RNA were found to be associated with stress granules in J1.1 cells treated with Nef exosomes compared to control exosomes (Fig. 3C).

The restriction factor APOBEC3G (A3G) is shown to bind HIV-1 mRNA and sequester it to stress granules [35]. We also found the levels of A3G to be upregulated in J1.1 cells treated with Nef exosomes (Fig. 4A), suggesting that increased levels of A3G might be responsible for sequestering viral mRNA into stress granules. Since A3G is also packaged in virus particles and reduces HIV-1 replication in the next round of infection, we checked if increased levels of A3G have any effect on the replicative ability of virions produced from J1.1 cells following their exposure to Nef exosomes. For this, we took equal amounts of p24 released from J1.1 cells treated with Nef or control exosomes, and checked their infectivity on TZMbl cells. Virions released from J1.1 cells treated with Nef exosomes were about three-fold less infectious than those released from cells treated with control exosomes (Fig. 4B).

**Figure 4:**
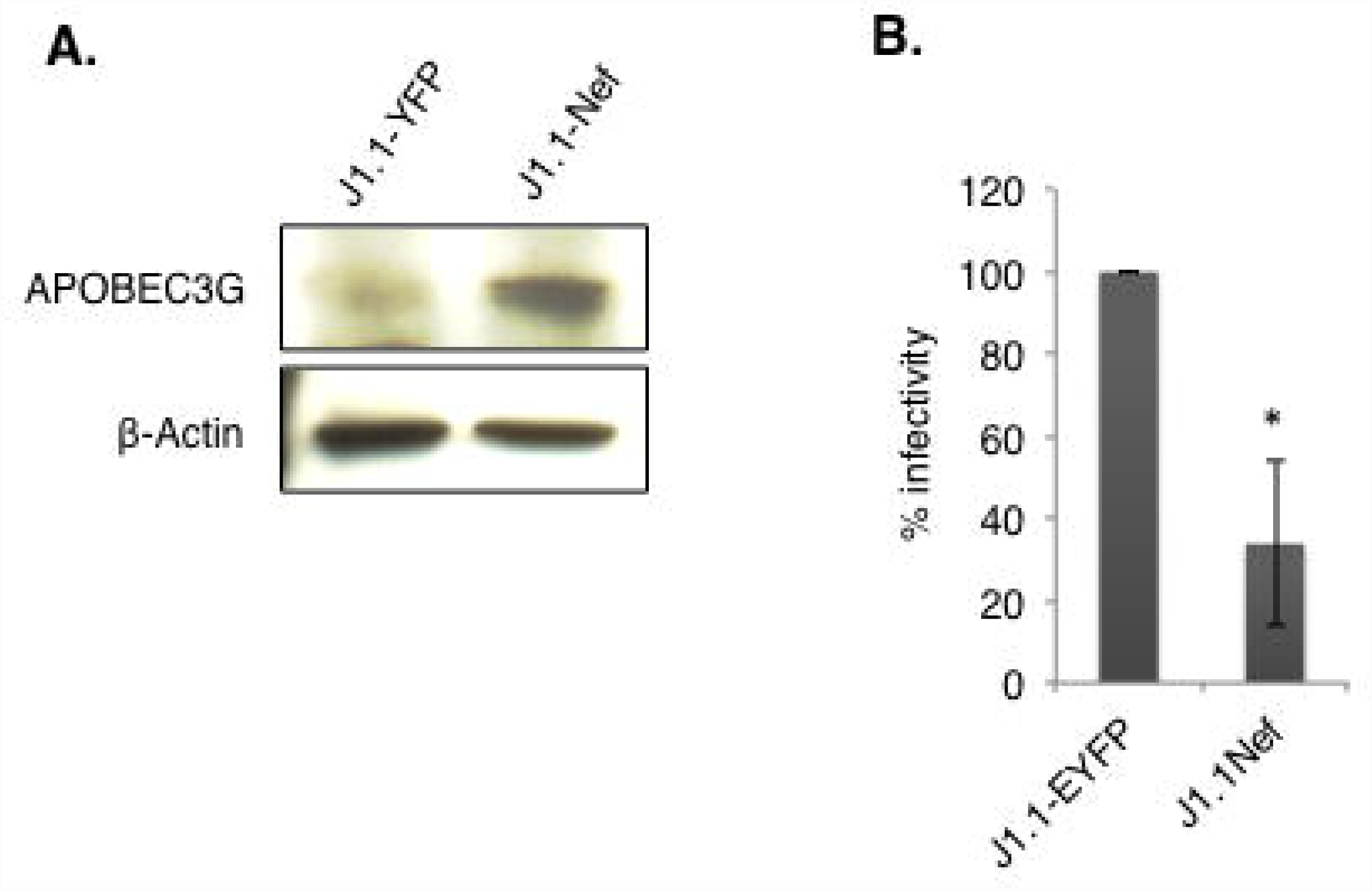
J1.1 cells treated with Nef exosomes produce less infectious virus. (A) J1.1 cells were treated with conditioned media from U937/EYFP cells (J1.1-YFP) or U937/Nef-EYFP cells (J1.1-Nef) for 72 hr. The cell lysates were then analyzed for APOBEC3G levels by Western blotting. Beta-actin served as a loading control. (B) Viruses in the culture media of J1.1 cells after treatment with conditioned media from U937/EYFP cells (J1.1-EYFP) or U937/Nef-EYFP cells (J1.1-Nef) for 72 hr, were assayed for infectivity on TZMbl cells and the values were normalized to p24 levels. Results are shown as mean of three independent experiments, each with triplicate measurements (*p<0.05).

## Discussion

Several reports have established the secretion of Nef into exosomes from HIV-1 infected T cells, monocytes and dendritic cells as well as from cells expressing Nef alone [25, 27, 28]. Nef alters membrane trafficking, increases the proliferation of MVBs and enhances its own secretion in exosomes [36-38]. An earlier study has shown that Nef secreted in exosomes causes activation-induced cell-death in bystander (uninfected) CD4+ T cells [28]. However, the effect of Nef exosomes on infected cells has not been studied. Here we analyzed the effects of Nef exosomes on the HIV-1 infected J1.1 CD4+ T cell line. Our results show that exosomal Nef reduces viral replication in infected cells, and that these effects are linked to the secretion of Nef into exosomes and its transfer to J1.1 cells. We further show that exosomal Nef does not affect proviral transcription or miRNA-mediated post-transcriptional silencing of viral mRNA, but results in increased assembly of stress granules in recipient cells, implying that Nef exosomes attenuate the translation of viral mRNAs in infected cells. We found decreased production of the viral Gag proteins (p55 and p24) from infected J1.1 cells upon co-culture with Nef-U937 cells or treatment with culture supernatant. The factor in culture supernatants responsible for reduced virus replication in infected cells was found to be exosomes. These nanovesicles have been reported to play important roles in intercellular communication by transporting proteins, mRNAs and miRNAs to recipient cells [39]. Exosomes produced from HIV-1 infected cells carry the entire viral genome, implicating their role in the spread of infection [40].

Stress granules are cytoplasmic bodies assembled in eukaryotic cells under different environmental stress conditions including oxidative stress, hypoxia and infection. The formation of stress granules leads to the translational arrest of select mRNAs and to enhance the translation of mRNAs that are involved in relieving stress [33]. Many reports have shown high levels of oxidative stress in patients with HIV-1 infection [41-44]. However, the cause of systemic oxidative stress in HIV-1 infected individuals has not been identified. We show that Nef exosomes induce the phosphorylation of eIF2α in J1.1 cells, which is a key signal for the assembly of stress granules that are sites of translational suppression of selected mRNAs [45]. Exosomal Nef attenuated virus production from infected cells not due to effects on HIV-1 transcription or miRNA-mediated translational repression, but due to the increased assembly of stress granules. In support of this, RNA immunopreicpitation showed increased association of gag mRNA with stress granules in infected cells that were exposed to Nef exosomes. The anti-HIV restriction factor, APOBEC3G selectively binds and shuttles HIV-1 mRNA between stress granules and polysomes [35]. We found increased levels of A3G in J1.1 cells treated with Nef exosomes and virions produced from these cells showed reduced infectivity.

## Conclusion

This study describes a novel function for Nef that is secreted in exosomes. The limitation of our data is that we cannot distinguish whether this is a direct effect of Nef or some other factor that is co-transported into exosomes with Nef. In either case, instead of the expected Nef-mediated activation of infected cells and increased viral replication, we found exosomal Nef to attenuate viral replication. This might be a viral strategy to limit the activation-induced death of infected cells, and thus aid in viral persistence.

## Author’s contribution

MYA performed most of the experiments, with help from HI and ZA. NK generated the U937 stable cell lines expressing mutant Nef proteins. MYA and SJ analyzed and interpreted the data, and wrote the manuscript.

## Acknowledgments

We thank Dr. Jomon Joseph at National Centre for Cell Sciences, Pune, India for the G3BP1-GFP plasmid.

